# Hits-Based Quantitative Characterization of SOBI-Recovered P3 Network Configuration: an EEG Source-Imaging Study

**DOI:** 10.1101/2020.09.24.312587

**Authors:** Adam John Privitera, Renee Fung, Yunqing Hua, Akaysha C. Tang

**Affiliations:** Neuroscience for Education Group, Faculty of Education, The University of Hong Kong, Hong Kong, China; Neural Dialogue Shenzhen Educational Technology & Neuroscience for Education Group, Faculty of Education, The University of Hong Kong, Hong Kong, China

**Keywords:** neuromarker, biomarker, EEG source imaging, source localization, BESA, LORETA, SOBI, ICA, P300

## Abstract

One frequently studied biomarker for health and disease conditions is the P3 component extracted from scalp recorded electroencephalography (EEG). The spatial origin of this significant neural signal is known to be distributed, typically involving large regions of the cerebral cortex as well as subcortical structures. Unlike the temporal characterization of the P3 by amplitude or latency measures from event-related potentials (ERPs), the spatial characterization of the P3 component is relatively rare, typically qualitative, and often reported as differences between populations (group differences between healthy controls and clinical groups). Here we introduce a novel approach to quantitatively characterize the spatial origin of the P3 component by (1) applying second-order blind identification (SOBI) to continuous, high-density EEG data to extract the P3 component, (2) modeling the underlying generators of the SOBI P3 component as a set of equivalent current dipoles (ECDs) in Talairach space using BESA; (3) using the application Talairach Client to determine the “hits” associated with the anatomical structures at three level of resolution (lobe, gyrus, and cell type). We show that the hits information provided by Talairach Client can enable a quantitative characterization of the spatial configuration of the network underlying the P3 component (P3N) via two quantities: cross-individual reliability (or consistency) of a given brain structure as a part of the P3N, and within-individual contribution of a given brain structure to the whole P3N network. We suggest that this method may be used to further differentiate individuals in the absence of differences in P3 amplitude or latency, or when scientific questions or practical application cannot be supported by a yes-no answer regarding the source of a P3 component. Finally, application of our method to a group of 13 participants revealed that frontal structures, particularly BA10, play a special role in the function of a global cortical network underlying novelty processing.

## I. Introduction

This manuscript describes a method for providing quantitative characterization of the spatial configuration of the neural network underling an extensively studied biomarker, the P300 (P3) [1-3], using EEG. Previously, information regarding neural generators underlying P3 components were initially coming from indirect evidence from non-scalp recorded EEG data, such as functional magnetic resonance imaging (fMRI) which measures the blood oxygen levels (BOLD) while an individual performs the same task during which the P3 component can be recorded with EEG [4], or using EEG recording made with implanted electrodes in patients who have these implants for medical reasons [5, 6]. More recently, P3 components have been localized with low resolution brain electromagnetic tomography (LORETA) which offers volume-source models [7, 8]. In contrast, point-source modeling using equivalent current dipoles (ECDs) has also been used to model the P3 components [9, 10]. All of these methods appear to support a general conclusion that P3 component reflects neural signals from a set of widely distributed brain structures. Here, we offer results that show how a hits-based analysis method can be used to address two new types of questions: how consistently, across different individuals, a particular neuroanatomical structure is involved the whole network that underlies the P3 components (P3N), and how much a given neuroanatomical structure within a given individual is contributing to the P3N.

## II. Methods

### A. Experimental procedures

Approval for this study was granted by the Human Research Ethics Committee of the University of Hong Kong. Thirteen normal functioning participants (6 males) between 19-33 years of age (M = 26.50 ± 4.48 years) performed a two-stimulus color visual oddball task during which the participants were instructed to quickly press the space button on a standard keyboard when a rare stimulus appeared (i.e. red rectangle) and not press the button when a frequent stimulus appeared (i.e. blue rectangle). Stimuli were presented for 500 ms with a variable inter-trial interval between 200-700 ms. In total, the task contained 250 trials (20% rare stimuli) and took approximately 15 minutes to complete.

Continuous reference-free EEG data were collected in an unshielded room using an active 128-channel Ag/AgCl electrode cap, ActiCHamp amplifier, and PyCorder data acquisition software (Brain Vision, LLC) with a sampling rate of 1000 Hz. Data collection began only after the impedance of all electrodes was below 10 KΩ. Before additional processing, a 50 Hz notch filter was applied to raw EEG data in order to remove line noise. Data were spatially down-sampled to 64-channel to allow for comparison with our previous work on localization [9].

### B. Extraction of the P3 network (P3N) using second-order bind source separation (SOBI)

SOBI [11] was applied to continuous EEG data to decompose the *n*-channel EEG data into *n*-components (**Fig. 1, Step 1**), each of which corresponds to a recovered putative source that contributes to the scalp recorded EEG signals. Detailed descriptions of SOBI’s usage [12-17], SOBI validation [18, 19], and review of SOBI usage [20-22] can be found elsewhere.

**Fig. 1.**
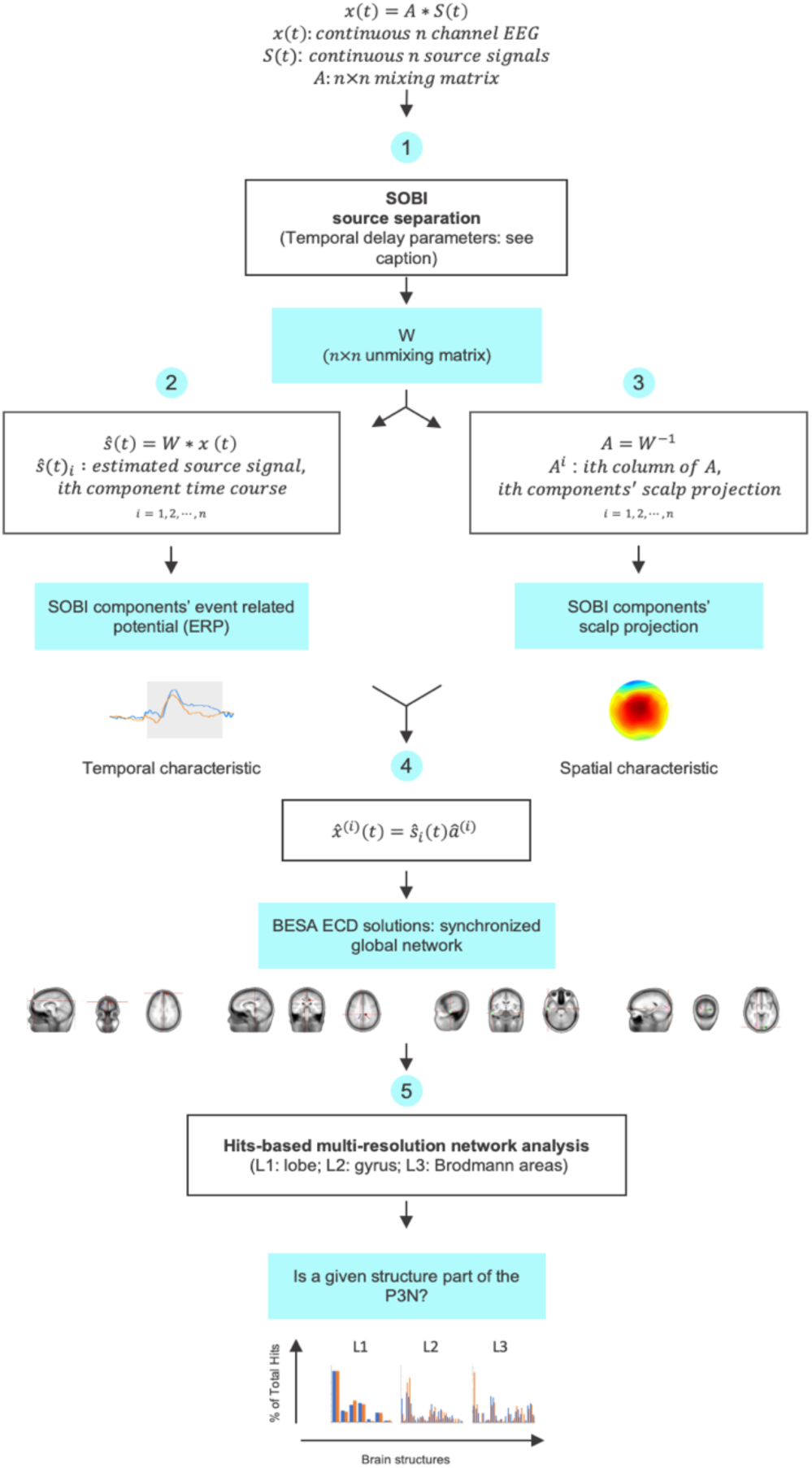
The analysis pipeline consists of five processing steps: (1) blind source separation using SOBI, (2 & 3) component selection, (4) equivalent current dipole modeling, and (5) quantitative multi-resolution network description. Orange and blue ERPs represent response to frequent and rare stimuli, respectively. The following set of delays (in ms) were selected based on our previous work using SOBI to recover components from EEG data.

Let x(*t*) represent the n continuous time series from the n EEG channels, where x_i_(*t*) corresponds to the ith EEG channel. Because various underlying sources are summed via volume conduction to give rise to the scalp EEG, each of the x_i_(*t*) is assumed to be an instantaneous linear mixture of *n* unknown components or sources s_*i*_(*t*), via an unknown *n*×*n* mixing matrix A,

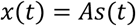

The putative sources, ŝ_i_(*t*) are given by

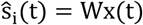

Where, *W*=*A*^−1^. SOBI finds the unmixing matrix *W* through an iterative process that minimizes the sum squared cross-correlations between one recovered component at time t and another at time t+τ, across a set of time delays. The following set of delays, τs (in ms), was chosen to cover a reasonably wide interval without extending beyond the support of the autocorrelation function:

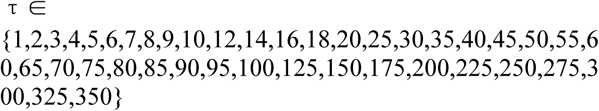

The ith component’s time course is given by ŝ_i_(*t*). The spatial location of the ith component is determined by the ith column of *A* (referred to as the component’s sensor weights or sensor space projection; **Fig 1, Step 3**), where *A = W*^−1^

Continuous EEG data were processed using SOBI [11]. **Fig. 1 Steps 1-3** show how spatial and temporal criterion are used to identify the SOBI P3 component. Components were identified as candidates for P3 if they displayed a single, symmetric, broadly-distributed dipolar field. Candidate components were further screened temporally for the presence of a large (i.e. amplitude greater than 2 μv) positive-going wave with a peak latency between 300-800 ms post stimulus. **Fig. 1 Step 4** shows the inputs used by BESA (BESA Research 6.1, Brain Electrical Source Analysis) to model the P3N. Equivalent current dipole (ECD) models were used to fit the scalp projection of P3 components.

Based on the nature of neuromodulatory innervation [23-25] along with previous localization work on the P3 [reviewed in 2], we anticipated that underlying generators would be broadly-distributed across the neocortex. Therefore, we began ECD model fitting with a fixed starting configuration, consisting of four pairs of bilaterally symmetrically placed dipoles at broadly-distributed locations **(Fig. 2)**. The following dipole locations, given in Talairach coordinates [26] were used as a starting template (X Y Z): ±20 63 24, ±51 9 - 28, ±44 −3 49, and ±24 −80 3.

**Fig. 2.**
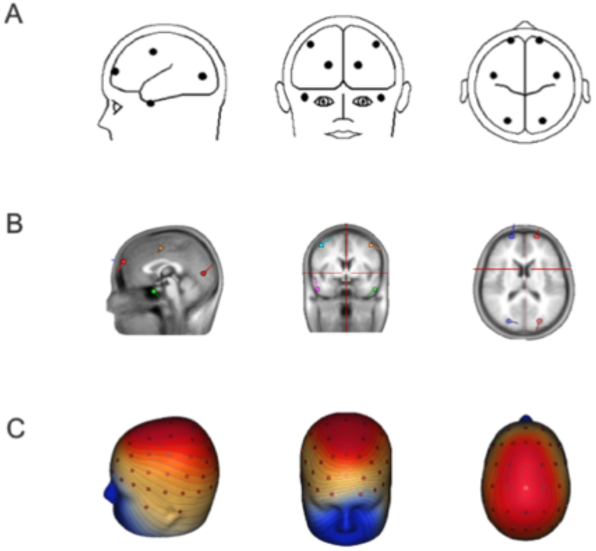
Model global network configuration used for source localization of P3N. **A**. ECD template consisting of 4 pairs of symmetrically placed dipoles used as the starting point for ECD modeling of the P3N in all participants. **B**. ECD template shown against the structural MRI (sMRI) of an average brain. **C**. 3D voltage maps of the ECD template model projection.

For efficiency, we first used a Four-Shell Ellipsoidal option for adults (CR=80) to fit the projected ERP wavefroms over a time window of 1000 ms (200 pre, 800 post-stimulus), then refitted with the Realistic Approximation option. Note that SOBI components’ scalp projections are time invariant (see equation in **Fig. 1, Step 3**), therefore, theoretically, the fitting of the model at any time point should make no difference in the final solution. However, due to numerical error in computation, fitting at different time points would produce non-identical final solutions. By using BESA’s option to fit over a large time window, one can improve stability and avoid an arbitrary decision to fit at one time point over another.

The resulting model is considered final if (1) the goodness of fit value is >=90% and, (2) the variations in voltage values in the residual map are within the range of background variations of the ERP waveform and, (3) removing any pairs of the ECDs will result in conditions (1) or (2) being unmet. Following this procedure, the resulting final models may have more or fewer pairs of ECDs than the initial starting network configuration.

### C. Analysis of hits associated with anatomical structures

BESA solutions in the form of Talairach coordinates were used as inputs to Talairach Client (version 2.4.3) [27] to identify anatomical structures contained within an 11 × 11 × 11 mm^3^ volume centered on each coordinate, and their respective hits numbers [27]. Hits number is a volumetric measure (unit: 1 mm^3^) reported for each structure in a Talairach Client solution. Because we were interested in identifying the underlying neural generators of the P3N, only data from grey matter structures were analyzed. A search range of ±5 mm (i.e. 11 × 11 × 11 mm^3^) was used to collect hits regardless of its distance to the center of the cube.

While it is common to take the nearest grey matter structure as the final solution, this choice can lead to structures occupying larger volumes being excluded from consideration, potentially resulting in an incomplete or inaccurate characterization of neural generators underlying a component. Instead, we perform further analysis on the full hits information and let the analysis drives the final solution. Therefore, in our solutions, each P3N is associated with a set hits-structure pairs at three levels of analysis.

Because the hits numbers vary a great deal both across-structure within an individual (within-participant variations in contribution) and across-individual for any given structure (cross-participant reliability), we provided two measures accordingly. The within-participant variability can be quantitatively indexed by differences in hits numbers associated with each structure in proportion to the total grey matter hits (% of total hits). The cross-participant reliability can be measured by the probability of a structure being observed across individuals, (i.e. % of participants that had at least 1 hit in that structure).

### D. Statistical analysis

Because hits numbers were not normally distributed, nonparametric statistics were used for all analyses. Binomial tests were performed on the proportion of participants with hits in a given region (1-tailed). One-sample Wilcoxon signed-rank tests (1-tailed) were performed on the % of total hits for each structure to test for a median significantly greater than 0. Significance levels were adjusted for multiple test when appropriate. All statistical analyses were performed using SPSS.

## III. Results

### A. Cross-participant variations in P3N configuration as a source of individual differences

An example of an extracted P3N at the level of an individual is shown in **Fig. 3A-D**. The 3D scalp projection of the SOBI P3 component (Data) closely matched the projection of the ECD solution (Model) with 99% goodness of fit (GoF) and little variance un-accounted (Residual) (**Fig. 3B)**. The number of dipole pairs needed to reach fitting criterion ranged between 3 and 5 indicating the existence of cross-individual differences in network configuration. The ECDs were widely distributed across all lobes of the neocortex (**Fig. 3C**), with no apparent focal clustering when ECDs from all participants were overlaid (**Fig. 3D**). This further indicates a high cross-participant variability in the precise configuration of the P3N. Thus, quantifying this cross-participant variability can offer previously unavailable ways to capture individual differences beyond those measured by temporal differences.

**Fig. 3.**
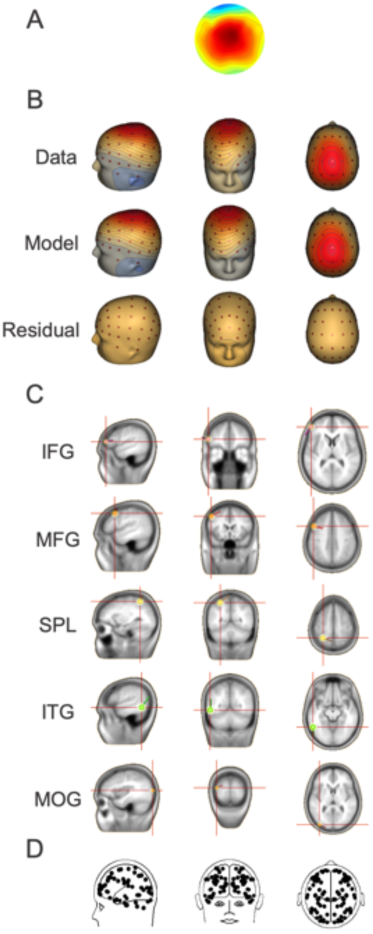
SOBI-recovered P3 Components (P3 Comps) and underlying network of generators (P3N). **A**. 2D scalp projection from individual participant. **B**. P3 scalp projection in voltage maps (data), ECD model projection (model), and the difference between data and model (residual). **C**. Each of the 5 pairs of ECDs shown against the structural MRI (sMRI) of an average brain. **D**. Overlay of dipoles from all participants’ P3Ns.

### B. Cross-individual reliability

There were 7 (lobe level), 33 (gyrus level), and 34 (cell type level) anatomical structures receiving at least 1 hit in Talairach Client’s output (subset of structures shown in **Table 1:A-C**, respectively). The number of hits (N-hits) associated with each brain structure varied greatly across individuals from zero to thousands at the lobe level and hundreds at the remaining two levels. The observation that the minimum N-hits were zero for most structures indicates that no structure was consistently observed across all individuals with the exception of the frontal lobe where the minimum N-hits was nearly 700. The average numbers of participants (N) in which a brain structure was observed were 8 at the lobe level, 5 at the gyrus level, and 5 at the cell type level, further confirming the considerable amount of *cross-individual variability*.

**Table 1.** N-Hits analysis of P3N across all participants for brain structures at the Lobe, Gyrus, and Cell Type Levels. N-hits: the number of hits associated with a structure for a given P3 Comp. % of total hits: N-hits in proportion of total grey matter hits associated with a P3 Comp. Only structures whose estimated contribution to the P3N was statistically significant are listed (one-sample Wilcoxon signed-rank test (1-tailed)).

| A Lobe Level |  |  |  |  |  |
| --- | --- | --- | --- | --- | --- |
| Structure | Range | M (SEM) | % of Total | Z | p |
| Frontal | 665 - 3451 | 2009 (215) | 45% | 3.180 | .001 |
| Parietal | 0 - 1829 | 470 (181) | 9% | 2.032 | .021 |
| Temporal | 0 - 1895 | 808 (149) | 19% | 2.937 | .002 |
| Occipital | 0 - 1318 | 684 (131) | 16% | 2.812 | .003 |
| Cerebellum | 0 - 3070 | 364 (245) | 8% | 1.826 | .034 |

| B Gyrus Level |  |  |  |  |  |
| --- | --- | --- | --- | --- | --- |
| Structure | Range | M (SEM) | % of Total | Z | p |
| IFG | 0 - 1187 | 192 (100) | 4% | 2.023 | .022 |
| MeFG | 0 - 327 | 51 (25) | 1% | 2.232 | .013 |
| MFG | 0 - 1789 | 809 (159) | 17% | 2.810 | .003 |
| SFG | 0 - 1655 | 786 (146) | 19% | 2.936 | .002 |
| PrG | 0 - 1017 | 170 (88) | 3% | 2.201 | .014 |
| SPL | 0 - 675 | 118 (61) | 2% | 1.826 | .034 |
| Pcun | 0 - 849 | 120 (68) | 2% | 2.032 | .021 |
| ITG | 0 - 576 | 201 (65) | 6% | 2.201 | .014 |
| MTG | 0 - 1653 | 180 (126) | 4% | 2.060 | .020 |
| STG | 0 - 1348 | 326 (146) | 7% | 2.032 | .021 |
| FuG | 0 - 404 | 86 (40) | 3% | 2.032 | .021 |
| MOG | 0 - 1298 | 261 (112) | 5% | 2.366 | .009 |
| LgG | 0 - 919 | 173 (91) | 5% | 1.841 | .033 |
| Cun | 0 - 1023 | 169 (84) | 4% | 2.201 | .014 |

Table. 1. N-Hits analysis of P3N across all participants for brain structures at the Lobe, Gyrus, and Cell Type Levels.
| C Cell Type Level |  |  |  |  |  |
| --- | --- | --- | --- | --- | --- |
| Structure | Range | M (SEM) | % of Total | Z | p |
| BA 10 | 0 - 1475 | 676 (518) | 22% | 2.805 | .003 |
| BA 9 | 0 - 934 | 319 (358) | 8% | 2.521 | .006 |
| BA 46 | 0 - 1218 | 188 (420) | 4% | 1.826 | .034 |
| BA 47 | 0 - 283 | 52 (97) | 1% | 1.841 | .033 |
| BA 8 | 0 - 1234 | 346 (479) | 7% | 2.207 | .014 |
| BA 6 | 0 - 1214 | 349 (501) | 9% | 2.201 | .014 |
| BA 7 | 0 - 985 | 193 (347) | 4% | 2.023 | .022 |
| BA 38 | 0 - 1348 | 312 (529) | 7% | 1.826 | .034 |
| BA 21 | 0 - 743 | 103 (216) | 2% | 1.826 | .034 |
| BA 20 | 0 - 882 | 202 (292) | 6% | 2.207 | .014 |
| BA 19 | 0 - 1196 | 278 (405) | 6% | 2.207 | .014 |
| BA 18 | 0 - 1019 | 345 (393) | 8% | 2.521 | .006 |
| BA 17 | 0 - 426 | 83 (152) | 3% | 1.826 | .034 |

At the lobe level, the probability of a structure being observed across individuals was significantly greater than chance for frontal (100%, p <=.001), temporal (85%, p=.011), and occipital lobes (77%, p=.046). These observations suggest that the P3N as captured by the SOBI P3 components would involve synchronized neural activity across at least the frontal, temporal, and occipital lobes, with the frontal lobe most reliably observed across individuals. Note that although no significant result was obtained for the parietal lobe in the present study, we believe that with increased power of the study, it is likely that the parietal lobe can also be found reliably with statistical significance. At the gyrus level, the SFG (85%, p=.011) and MFG (77%, p=.046) were observed across individuals with statistical significance. This is consistent with the observation at the lobe level that the frontal lobe has the highest cross-participant reliability. At the cell type level, only BA10 (85%, p=.011), a frontal structure known to play a special role in executive function [28], was found reliably statistically. The remaining structures at the gyrus and cell type levels had a probability of being observed of about 30% or below and did not reach statistical significance.

These cross-participant reliability measures suggest that the P3N appears to consist of two types of structures across individuals: reliably observed frontal structures, and highly variable non-frontal structures. This contrasting pattern of reliability and variability may point to an additional source of individual difference—that is the ability of the frontal lobe to synchronize, hence, engage with other more variably involved brain structures.

### C. Quantifying cross-individual variability in network configuration

In providing quantitative characterization of the spatial configuration of the P3N, the relative volumetric contribution of a given structure towards the whole network is operationally defined as the % of total hits. The more hits a structure has in proportion to the total hits associated with the P3N, the greater we consider its contribution to the whole P3N. **Table 1** showed that 5 structures at the lobe level, 14 at the gyrus level, and 13 at the cell type level made statistically significant contributions to the whole P3N (Wilcoxon signed-rank tests, 1-tailed). The % of total hits measures are highest for the frontal lobe structures (the frontal lobe: 45%; SFG: 19%, MFG: 17%; BA10: 22%). These frontal structures appear to be sufficiently deviated from the rest of the structures as shown in the histograms of % total hits measure (**Fig. 4**). Together, these analyses of cross-structure differences in their involvement in the P3N offer further support for a rather unique role played by frontal lobe structures, particularly BA10, in this global network.

**Fig. 4.**
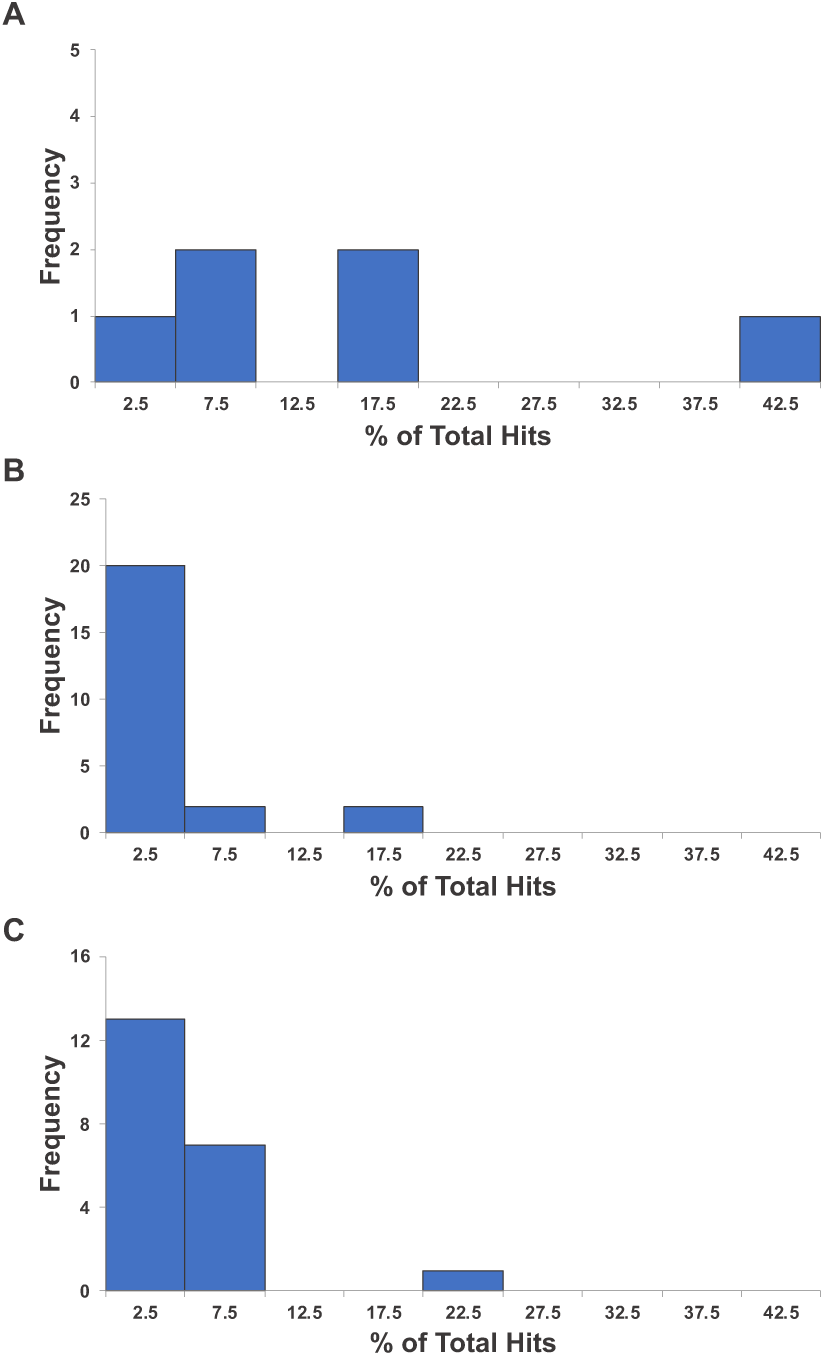
Histogram of % of total hits at Lobe (**A**), Gyrus (**B**), and Cell Type (**C**) levels. Average % of total hits at the lobe, gyrus, and cell type level were 16%, 4%, and 5%, respectively.

From the % of total hits measure, one can also determine the size of the P3N. The average P3N sizes (i.e. number of structures with N-hits >0) were 4 at the lobe level, 10 at the gyrus level, and 9 at the cell type level, with ranges being 3-6, 7-15, and 3-15 structures, respectively. Note that one must be cautious in interpreting these numbers because network size can be high if ECDs happen to be located in the area bordering multiple structures. Therefore, no further interpretation will be made on this measure here.

## IV. Conclusion

We presented results on the analysis of spatial configuration of the P3 network via a novel hits-based quantitative characterization of spatial configuration of P3N. Methodologically, we show that by analyzing the number of hits associated with anatomical structures at lobe, gyrus, and cell type-levels, both the contribution of constituent structures within the P3N spatial configuration and the consistency of each structure’s involvement across individuals can be quantified at multiple levels. Scientifically, we found that frontal lobe structures, particularly BA10, are reliably involved across individuals and make the greatest volumetric contribution to the P3N in comparison to other less reliably involved non-frontal structures. These variations revealed by hits analysis point to a previously little explored spatially-based source of individual differences.

## Acknowledgment

This work was supported by a grant from the University of Hong Kong [104004683] and a donation from the Sweeting Memorial Fund awarded to A.C.T. We thank Drs. R. Sun and E. Tsang for their assistance.

